# C-terminal domain of SARS-CoV-2 nucleocapsid protein binds transcriptional regulatory sequence RNA with high affinity and forms higher-order assemblies

**DOI:** 10.64898/2026.10.02.756193

**Authors:** Parul Gupta, Mahavir Singh

**Affiliations:** Molecular Biophysics Unit, Indian Institute of Science, Bengaluru, 560012, INDIA

**Author notes:** **Corresponding Author:** Mahavir Singh - Molecular Biophysics Unit, Indian Institute of Science, Bengaluru, 560012, INDIA. **Present Addresses:** Parul Gupta - Molecular Biophysics Unit, Indian Institute of Science, Bengaluru, 560012, INDIA.

**Keywords:** SARS-CoV-2, nucleocapsid, TRS, NMR, ITC

## Abstract

The SARS-CoV-2 nucleocapsid (N) protein is a 419-amino-acid modular protein essential for viral RNA replication, genome packaging, and host-cell modulation. It contains two structured domains, the N-terminal RNA-binding domain (NTD) and C-terminal dimerization domain (CTD), separated and flanked by three intrinsically disordered regions. Although the CTD mediates N-protein dimerization and contributes to RNA binding, the molecular basis of its interaction with RNA remains poorly understood. Here, we characterized CTD binding to transcriptional regulatory sequence (TRS) RNAs from the SARS-CoV-2 genome using NMR spectroscopy, isothermal titration calorimetry, size-exclusion chromatography, and in vitro phase-separation assays. The CTD bound structured stem-loop TRS RNA with higher affinity than the canonical RNA-binding NTD. NMR chemical-shift perturbation experiments indicated formation of higher-order molecular assemblies upon titration of the CTD with longer, structured stem-loop TRS RNA, consistent with size-exclusion chromatography results. The CTD also underwent RNA-induced phase separation *in vitro*, with stem-loop TRS RNA promoting markedly more robust condensation than single-stranded RNA sequences. Together, these findings establish the CTD as a high-affinity, structure-selective RNA-binding domain with a preference for stem-loop TRS RNA. The results further show that CTD-TRS interactions promote higher-order assembly and phase separation, providing mechanistic insight into N-protein-mediated viral genome packaging and nucleocapsid assembly.

## Introduction

Severe acute respiratory syndrome coronavirus 2 (SARS-CoV-2) is a member of the coronavirus (CoV) family that caused the COVID-19 pandemic. Coronaviruses (CoVs) are enveloped positive-sense single-stranded RNA viruses that have emerged as significant human pathogens over the past five decades ^1–3^. The genome of SARS-CoV-2 virus is ∼30 kilo-bases (kb) long which is one of the largest known RNA genomes ^1,4,5^. The virus requires efficient interplay of viral proteins, viral genome, viral RNA-binding proteins and host proteins to perform replication, transcription and packaging of virus ^6–9^.

SARS-CoV-2 encodes four structural proteins: spike (S), envelope (E), membrane (M), and nucleocapsid (N), of which the N protein is most abundantly expressed during infection ^10–12^. Beyond its canonical role in packaging the viral genome to form the ribonucleoprotein (RNP) complex, the N protein performs diverse functions, including regulation of viral RNA synthesis, modulation of host innate immune responses, and interaction with numerous host RNA-binding and stress granule-associated proteins ^13–18^. The SARS-CoV-2 N protein consists of an N-terminal intrinsically disordered region (IDR) (N-arm), a folded N-terminal RNA-binding domain (NTD), a serine/arginine-rich linker region (Lkr), a folded C-terminal dimerization domain (CTD), and a C-terminal intrinsically disordered region (C-tail) (Figure 1A). While the NTD is widely regarded as the primary RNA-binding domain (Figure 1B), increasing number of evidences suggest that multiple regions of N protein, beside the NTD contribute to RNA recognition in a cooperative and context-dependent manner ^13,19–23^.

**Figure 1.**
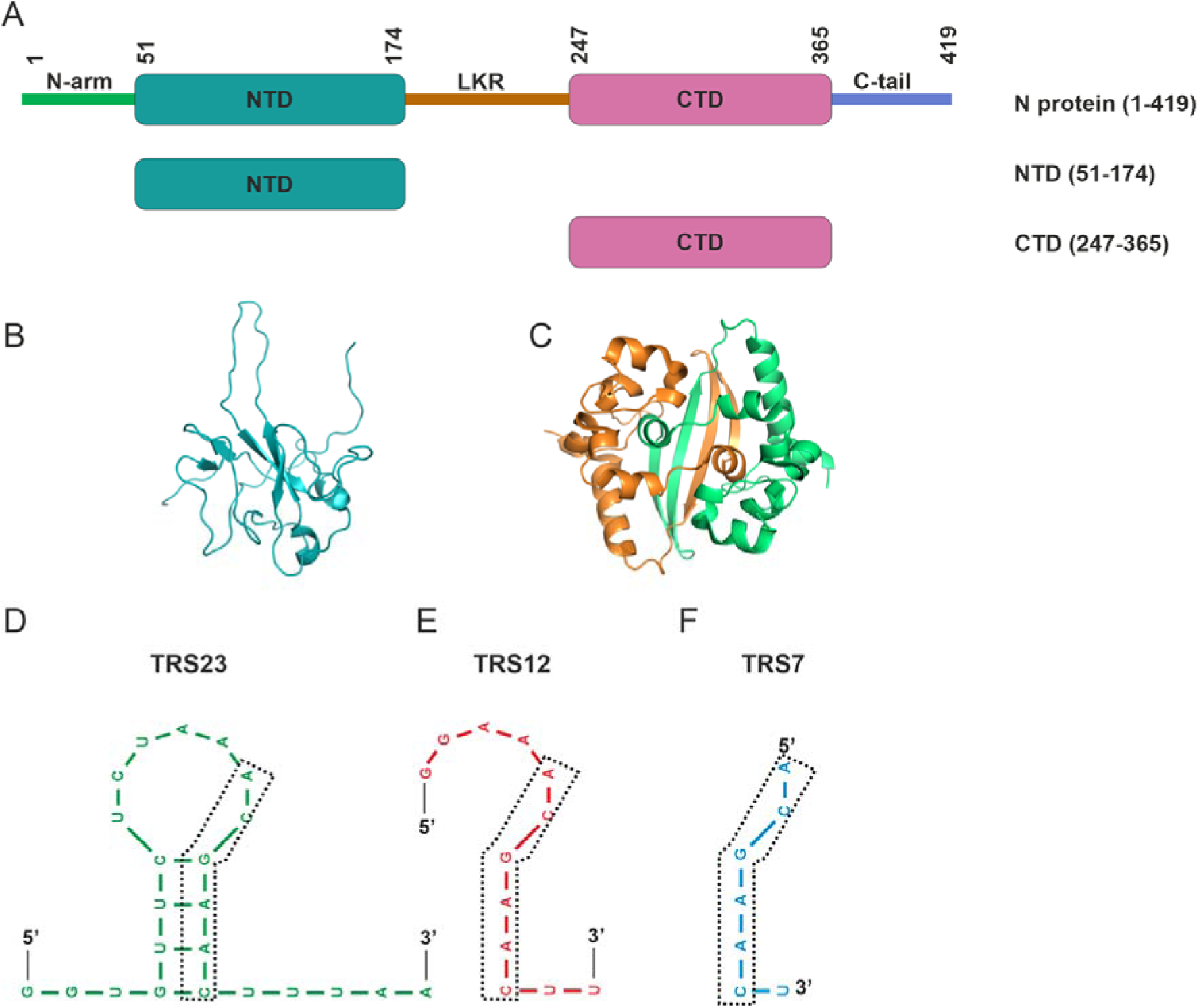
Domains of SARS-CoV-2 nucleocapsid (N) protein and transcription-regulatory sequences (TRS) RNA used in this study. **(A)** Schematic representation of the full-length N protein (residues 1–419). The N protein consists of an N-terminal arm (N-arm, green), the N-terminal domain (NTD, cyan, residues 51–174), a central linker region (Lkr, orange), the C-terminal domain (CTD, magenta, residues 247–365), and a C-terminal tail (C-tail, blue). Folded domains used in this study are shown with their domain boundaries: NTD and CTD. **(B)** Cartoon representation of the NMR solution structure of NTD (PDB: 6YI3, cyan). **(C)** Cartoon representation of the crystal structure of CTD homodimer (PDB: 6YUN). The two protomers of the CTD dimer are shown in different colours (orange and green). **(D–F)** Mfold predicted structures of representative transcription-regulatory sequences (TRS) from the SARS-CoV-2 genome used in this study: TRS23 **(D)**, TRS12 **(E)**, and TRS7 **(F)**. Black box represents the core TRS sequence.

One of the characteristic features of coronaviruses is the gene expression by discontinuous transcription ^9^. This generates a nested set of subgenomic mRNA (sgmRNAs). During negative-strand (–) RNA synthesis, the viral RNA-dependent RNA polymerase (RdRp) undergoes template switching guided by conserved transcription regulatory sequences (TRSs) to generate sub-genomic mRNAs (sgmRNAs). These TRSs are located in the leader region (TRS-L) of 5’ untranslated region (UTR) and upstream of each open reading frame in the body of the genome (TRS-B) ^24,25^. Base pairing between TRS-B and TRS-L during (–) strand synthesis enables the fusion of the leader sequence with downstream gene segments, producing discontinuous (–) strand templates that are subsequently transcribed into sgmRNAs.

A previous transcriptomics analysis had revealed that SARS-CoV-2 infected cells generate not only canonical genomic RNA and nine major sgRNAs but also numerous non-canonical transcripts arising from alternative template-switching events ^26,27^. Among these, the nucleocapsid-encoding sgRNA is the most abundant, revealing the importance of N protein expression during infection. Zuniga et al. showed the impaired synthesis of sgmRNA on deletion of N protein coding region ^28^. However, this deletion did not affect the replication of genomic RNA, suggesting the role of N protein in discontinuous transcription ^28^. Moreover, phosphorylation-dependent recruitment of the host RNA helicase DDX1 by N protein has been shown to regulate the transition between discontinuous and continuous transcription modes ^29^. Despite these observations, the molecular mechanisms by which N protein interacts with TRS RNA during transcription remain poorly understood.

The CTD of N protein is primarily known as dimerization domain responsible for the N protein oligomerisation (Figure 1C). Previous structural study revealed that CTD forms stable dimers in solution and higher-order oligomers ^23,30^. Disruption of CTD-mediated self-association attenuates coronavirus replication, highlighting CTD oligomerization as a potential antiviral target ^23^. Importantly, surface electrostatic analyses of SARS-CoV-2 N-CTD have identified conserved positively charged surface along the α-helical interface of the dimer, similar to those observed in SARS-CoV and MERS-CoV, suggesting its potential role in nucleic acid binding.

Indeed, there are increasing number of studies that showed the direct interaction of RNA with CTD ^22,23,27,31–38^. NMR titration studies demonstrated that SARS-CoV-2 N-CTD binds the structured stem-loop II motif (s2m) RNA, inducing extensive line broadening consistent with intermediate-exchange binding while preserving CTD folding and dimerization ^39^. Two ATP binding pockets on the CTD dimer interface were identified and ATP was shown to compete with RNA to bind to CTD ^39^. Another study showed that CTD alone binds different constructs of TRS RNA with relatively weak affinity, whereas the CTD when connected with either IDRs (Lkr or C-tail) shows enhanced RNA binding ^27^. These studies also showed that unpaired adenosine dinucleotides at the 5′ end of the TRS-L stem-loop play a key role in nucleocapsid recognition, pointing to a structural basis for selective TRS engagement during transcription. Together, these findings establish the CTD as a critical domain playing role in RNA binding, N protein oligomerization, and host protein interactions.

Despite growing recognition of CTD’s involvement in RNA binding and transcriptional regulation, the molecular details governing CTD interactions with transcription regulatory sequence remain unclear. In particular, molecular details of CTD binding to TRS RNA, which is central elements in discontinuous transcription, has been lacking. Given the emerging multifunctionality of the CTD in N protein functions, elucidating the CTD-RNA interaction is important for a mechanistic understanding of coronavirus genome transcription and assembly.

In this study, we investigated the interaction between the SARS-CoV-2 N CTD and an TRS RNA sequences using biophysical approaches (Figure 1D-F). Our results showed that CTD binds to the stem-loop RNA with higher affinity than the single-stranded RNA (ssRNA). NMR titrations studies showed that the interaction of TRS RNAs with CTD is in the intermediate exchange in NMR timescale resulting in line broadening and disappearance of peaks. Size-exclusion chromatography studies confirmed the formation of oligomeric complexes on interaction of stem-loop RNA with the CTD. Furthermore, we showed that CTD can undergo *in vitro* phase separation in the presence of RNA, wherein the presence of stem-loop TRS RNA induced robust phase separation of the CTD. Taken together, these results show that the CTD of N protein uses its intrinsic property of dimerization and RNA binding to form higher oligomeric RNA-protein complexes, which may have implication for viral nucleocapsid assembly.

## Materials and Methods

### Protein expression & purification

The gene encoding the SARS-CoV-2 N protein C-terminal dimerisation domain (CTD), fused to an N-terminal hexa-histidine tag, was codon-optimised and synthesised by GenScript and cloned into the pET-28a(+) expression vector. The plasmid was transformed into *E. coli* BL21(DE3) competent cells, and grown overnight at 37°C, 180 rpm, followed by the secondary culture at 37°C, 180 rpm till OD_600_ ∼0x.8. The cells were induced with 250 µM IPTG after the optical density of ∼0.8 at 600 nm, followed by incubation at 18 °C for 16 h to promote soluble expression. Cells were harvested by centrifugation (5000 rpm, 15 min, 4 °C), and the resulting pellet was resuspended in lysis buffer containing 25 mM Tris-HCl, pH 7.4, 500 mM NaCl, 2 mM MgCl₂, 5 mM β-mercaptoethanol and 5 mM imidazole. Cells were lysed by sonication on ice. The lysate was clarified by centrifugation at 13,000 rpm for 1 h at 4 °C, and the supernatant was loaded onto a pre-equilibrated Ni-NTA affinity column. The column was washed with wash buffer (lysis buffer + 20 mM imidazole) to remove non-specifically bound proteins. The bound CTD was eluted using the elution buffer (lysis buffer + 400 mM imidazole). Eluted fractions containing the target protein were pooled and concentrated using a centrifugal concentrator (10 kDa MWCO). For final purification, the concentrated sample was loaded onto a Superdex 200 Increase size-exclusion chromatography (SEC) column, which was equilibrated with SEC buffer (20 mM sodium phosphate, 100 mM NaCl, pH 6.5). The major peak corresponding to dimeric CTD was collected, analysed by SDS-PAGE to confirm purity, and used for subsequent experiments. The concentrations of the protein were calculated by ultraviolet (UV) absorbance at 280 nm using a UV spectrophotometer (Eppendorf), and the calculated molar extinction coefficients of N-terminally hexa-histidine tagged CTD (□ = 16.96 mM^−1^ cm^−1^), determined using EXPASY webserver.

### RNA preparation

The TRS23 and TRS12 RNA sequences (Supplementary Table T1) were synthesised by *in vitro* transcription using specific DNA templates and T7 RNA polymerase using established protocols. After *in vitro* transcription, the RNA was precipitated with ethanol, and the resulting pellets were dissolved in Milli-Q water. The RNA was then purified through denaturing urea-TBE polyacrylamide gel electrophoresis. The bands corresponding to the desired products were visualised under UV light, excised from the gel, and recovered using a standard crush-and-soak method. To further enhance purity, the recovered RNAs were processed by anion-exchange chromatography on a 5 mL HiTrap Q HP column. The purified material was subsequently buffer exchanged into water using a centricon concentrator (Millipore). Before use, the RNAs were thermally annealed by heating to 95°C for 3 minutes and then rapidly cooled on ice for 25 minutes. TRS23 and TRS12 RNA constructs were designed to contain a 5′-terminal GG dinucleotide to enhance transcription efficiency with T7 polymerase.

The concentrations of the RNA were calculated by ultraviolet (UV) absorbance at 260 nm using a UV spectrophotometer (Eppendorf), and the calculated molar extinction coefficients of TRS23 (□ = 235.1 mM^−1^ cm^−1^), TRS12 (□ = 125.9 mM^−1^ cm^−1^), and TRS7 (□ = 71.1 mM^−1^ cm^−1^), determined using EXPASY webserver ^40^.

TRS7 RNA (Supplementary Table T1) was ordered commercially. RNA sample was HPLC purified and dissolved in appropriate buffer before use.

### NMR sample preparation, data acquisition, and processing

All NMR spectra (1D ^1^H and 2D ^1^H–^15^N HSQC) were acquired on a Bruker 700 MHz or 800 MHz spectrometer equipped with a cryogenic probe or a room temperature probe. The NMR spectra were processed and analysed using Bruker TopSpin or NMRpipe, and analysed using the NMRFAM-SPARKY software ^41,42^. The NMR buffer used was 20 mM NaH_2_PO_4_, pH 6.5, 100 mM NaCl. 10% D_2_O (v/v) (Cambridge Isotope Laboratories; Cat. No. DLM-4–25) was added to the sample for the spectrometer deuterium lock. Typically, for the titration experiments, protein sample concentration was kept between 100 - 200 μM (protein concentrations here refer to CTD monomer). Protein:RNA molar ratios quoted throughout are therefore CTD monomer:RNA. For example, a CTD:RNA ratio of 1:0.25 corresponds to 0.25 RNA molecules per CTD monomer, or 0.5 RNA molecules per CTD dimer.

1D-^1^H 1,1-echo NMR spectra of the imino region of TRS23 RNA were recorded. Uniformly ^15^N labelled CTD protein samples were titrated with different TRS RNA sequences, and a 2D ^1^H - ^15^N HSQC spectra were recorded at each step of titration.

For CTD-TRS23 and CTD-TRS12 titrations, the decrease in peak intensities of residues at each titration point was measured and peak intensity ratios (*I/I_0_*), where *I* and *I_0_* represent the peak intensities in the presence and absence of RNA, respectively was calculated. The intensity plots (*I/I_0_*vs residues) were plotted for each titration point.

The chemical shift perturbations (CSPs), for the CTD-TRS7 titration were analysed as combined amide chemical shift changes with the equation:

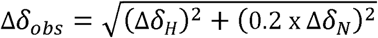

where the chemical shift changes (in ppm) between the free and RNA-bound states in the 1^H^ and 15^N^ dimensions are denoted by Δδ_H_ and Δδ_N_, respectively. The assignment of the CTD protein was taken from a previously published study ^43^, where the assignment of the CTD protein was deposited as BMRB entry 50518.

### Fitting and Simulating NMR Titration Profiles

^1^H - ^15^N HSQC NMR titration spectra of CTD and CTD:TRS7 were analysed using TITAN (TITration ANalysis) software ^44^. TITAN numerically simulates the evolution of magnetization during a pulse sequence in the presence of chemical exchange. Residues that showed a visible chemical shift perturbation upon TRS7 RNA binding, and whose peaks were not severely overlapped with neighbouring resonances, were selected for lineshape analysis. The K_d_ and K_off_ values were obtained by fitting experimental spectra to simulated spectra generated using a two-state binding model. Best-fit parameters were determined by minimizing the difference between experimental and simulated data. The standard deviation obtained from the mean of 100 bootstrap analysis was used to determine the errors in fitted parameters.

### Isothermal titration calorimetry (ITC)

We performed isothermal titration calorimetry (ITC) experiments to quantitate the binding of NTD and CTD with different RNAs. ITC experiments allows determination of thermodynamic parameters such as change in enthalpy (Δ*H*), change in entropy (Δ*S*), Gibbs free energy change (Δ*G*), equilibrium dissociation constant (*K*_d_), and stoichiometry (*n*) of interaction under a set of experimental conditions.

ITC experiments were performed using the PEAQ-ITC. CTD protein and RNAs were prepared in the same buffer consisting of 20 mM HEPES, pH 7.4, 150 mM NaCl. All the samples and buffer were thoroughly degassed before the experiment. The RNA and protein were quantified by measuring UV absorption at 260 nm and 280 nm wavelengths, respectively. For the titration experiments, the sample cell was filled with 20 μM RNA and titrated with 400 μM CTD protein (monomer concentration; equivalent to 200 μM CTD dimer) in a syringe. Nineteen injections of the titrant were performed with an interval of 180 s between each injection and a stirring speed of 800 rpm. The heat of dilution was subtracted from the integrated heat data, and the data were then fitted for the one-site binding model using Malvern PEAQ-ITC Analysis software provided by the vendor. All the parameters were kept floating during the data fitting. Experiments were repeated 2–3 times for consistency and reproducibility.

### AlphaFold3

AlphaFold 3 (AF3) predictions were used to generate the structural models of CTD in complex with TRS7, TRS12, and TRS23 RNAs, using the available Web server https://alphafoldserver.com ^45^. The FASTA sequence of CTD and RNA sequences were provided as input for AF3 modelling. For each CTD–RNA complex, multiple models were generated, and the highest-ranked model (based on the pTM and ipTM confidence scores > 0.8 was selected for further analysis. A 2:1 CTD:RNA stoichiometry was used for modelling the CTD-TRS23, whereas a 1:1 stoichiometry was used for the CTD-TRS12 and CTD-TRS7 complexes. The predicted complexes were visualized and analysed in PyMOL (Version 3.0 Schrödinger, LLC).

### *In vitro* phase separation

CTD was diluted to a final concentration of 100 µM, in phase separation buffer consisting of 20 mM HEPES (pH 7.4) and 100 mM NaCl. For droplet visualization, CTD was mixed with RNA at varying concentrations and gently pipetted to ensure homogeneity. Samples were imaged immediately after mixing the protein and RNA. Approximately 5 µL of each sample was transferred onto a glass slide and visualized using a Leica SP8 FALCON confocal microscope. Image acquisition parameters were kept constant for all experiments. Image analysis was performed using ImageJ (Fiji) ^46^.

## Results

### Interaction of NTD and CTD of N protein with the transcriptional regulatory sequence (TRS) RNA

The NTD and CTD constructs of the N protein were purified according to the protocol described in the Materials and Methods section. Size-exclusion chromatography profile showed that the NTD purifies as a monomer in solution. On the other hand, as expected, the CTD purifies as a dimer in solution (Supplementary Figure S1A and S1B). SDS–PAGE analysis was carried out to reveal the relative purity of the samples (Supplementary Figure S1C and S1D). The ^15^N-labeled samples of both the NTD and CTD were prepared and 2-dimensional (2D) ^1^H–^15^N HSQC spectra were recorded. The spectra showed well-dispersed resonance peaks for both NTD and CTD, indicative of well-folded proteins in solution (Supplementary Figure S1E and S1F).

A 23-nucleotide transcription regulatory sequence RNA (henceforth called TRS23) was selected for the initial interaction studies with the NTD and CTD of the N protein (Figure 1D). Secondary structure prediction using Mfold indicated that TRS23 adopts a stem–loop conformation containing four base-paired stem region as in the viral genome ^47^. The conserved, core TRS sequence (ACGAAC) remains as a part of the stem in TRS23 stem-loop RNA. TRS23 RNA was *in vitro* transcribed and purified as described in the Materials and Methods section. In addition, two variants of TRS RNA, a 12-nucleotide long TRS12 and 7 nucleotide long TRS7 were also prepared (Figure 1E and 1F). While TRS12 RNA was also prepared using *in vitro* transcription, TRS7 RNA was commercially synthesized (Sigma-aldrich). Both TRS12 and TRS7 contain core TRS sequence.

ITC experiments were performed to determine the binding of TRS RNAs to the NTD and CTD of N protein. In all the experiments, RNA sample was placed inside the sample cell and titrated with the protein (NTD or CTD) in the syringe. Figure 2 shows the representative raw and fitted binding isotherms, and Table 1 summarises the thermodynamic parameters obtained for each titration.

**Figure 2.**
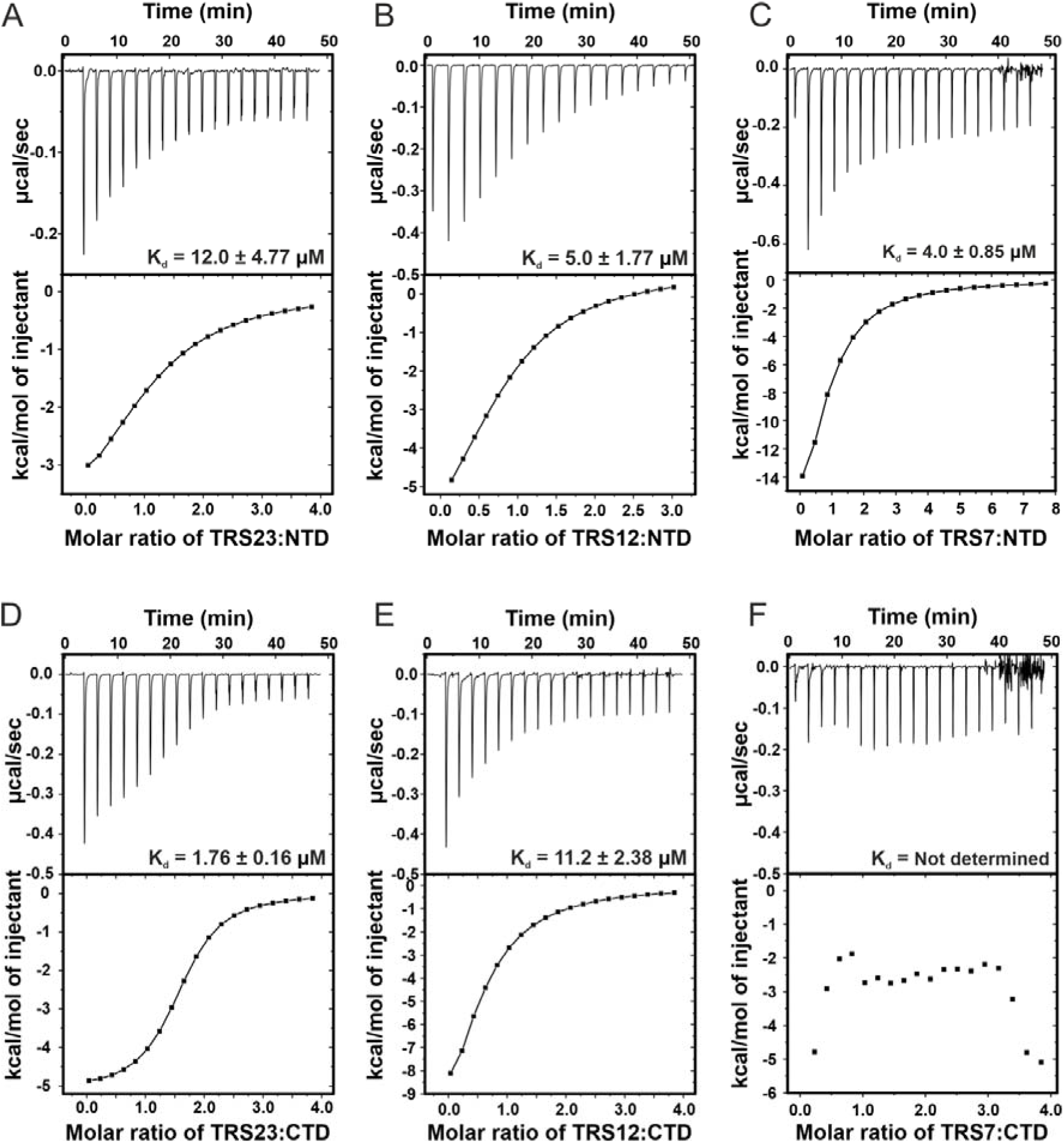
Binding of N protein’s CTD and NTD with TRS RNAs monitored using isothermal titration calorimetry. (**A**) Interaction of NTD with stem loop RNA TRS23. (**B**) Interaction of NTD with single stranded RNA TRS12. (**C**) Interaction of NTD with single stranded RNA TRS7. (**D**) Interaction of CTD with stem loop RNA TRS23. (**E**) Interaction of CTD with single stranded RNA TRS12. (**F**) Interaction of CTD with single stranded RNA TRS7. The top portion of each panel shows the raw titration data, and the bottom panel shows the integrated heats fitted to single-site binding isotherm. The equilibrium dissociation constants (K_d_) obtained upon fitting of the raw data is mentioned in each panel. No consistent heat change was observed for interaction between TRS7 and CTD.

**Table 1.** Equilibrium dissociation constants (K_d_s) and other thermodynamic parameters obtained from the interaction of NTD and CTD with different RNAs using ITC experiments at (errors are fitting errors from the reported data sets). Note that the CTD concentrations were determined on a monomer basis and n therefore reports CTD monomers bound per RNA molecule. NTD is monomeric in solution.

| S. No. | Experiment | $K_d$ (μM) | $\Delta G$<br>(kcal/mol) | $\Delta H$ (kcal/mol) | $-T\Delta S$<br>(kcal/mol) | N |
| --- | --- | --- | --- | --- | --- | --- |
| 1 | CTD-TRS23 | $1.76 \pm 0.16$ | -7.86 | $-5.13 \pm 0.10$ | -2.73 | $2.09 \pm 0.019$ |
| 2 | CTD-TRS12 | $11.2 \pm 2.38$ | -6.83 | $-10.6 \pm 3.82$ | 3.79 | $0.96 \pm 0.21$ |
| 3 | CTD-TRS7 | $34.5 \pm 4.67$ | - | - | - | - |
| 4 | NTD-TRS23 | $12.0 \pm 4.77$ | -6.72 | $-5.43 \pm 0.32$ | -1.29 | $0.89 \pm 0.14$ |
| 5 | NTD-TRS12 | $5.0 \pm 1.77$ | -6.44 | $-4.34 \pm 0.32$ | -2.1 | $0.9 \pm 0.3$ |
| 6 | NTD-TRS7 | $4.01 \pm 0.85$ | -7.37 | $-31.4 \pm 8.21$ | 24.1 | $0.8 \pm 0.13$ |

Binding of NTD to the stem–loop TRS23 RNA yielded a dissociation constant (K_d_) of approximately 12 μM, whereas interaction with the single-stranded TRS12 and TRS7 RNA resulted in a stronger affinities with the similar K_d_S of ∼5 μM and ∼4 μM respectively (Figure 2A-C) (Table 1). These observations suggest that the NTD preferentially binds single-stranded RNA (TRS12 and TRS7) over structured stem–loop RNA (TRS23).

In the next set of ITC experiment, three different RNAs were titrated with the CTD (Figure 2D-F). Titration of CTD into stem-loop TRS23 resulted in a stronger interaction with a K_d_ of ∼ 1.76 μM (Figure 2D. On the other hand CTD binds TRS12 RNA with a weaker affinity (K_d_ of ∼ 11.2 μM) (Figure 2E). In case of TRS7 RNA and CTD interaction, in the ITC experiment, we observed heat change of small magnitude, suggesting binding, however we could not reliably fit the data to extract a meaningful K_d_ (Figure 2F). This suggest a weak binding between CTD and TRS7 RNA, which was later confirmed by NMR based titrations that revealed a K_d_ of ∼ 34.5 μM (Table 1) (discussed later). These results confirmed that CTD can binds RNA with the preference for structured stem-loop RNA than the single stranded RNAs. Taken together, these results suggest distinct RNA recognition modes and affinities for the NTD and CTD of the N protein.

The fitted stoichiometries provide additional information about the binding modes. For the CTD-TRS23 titration, n = 2.09 ± 0.019 (Table 1). Since CTD was quantified as a monomer, this corresponds to two CTD monomers ( or one CTD dimer) is bound per TRS23 RNA molecule. In contrast, the CTD-TRS12 titration gave n = 0.96 ± 0.21, corresponding to one CTD monomer per TRS12 (or two TRS12 molecules bound per CTD dimer). The structured stem-loop TRS23 is therefore engaged by a single CTD dimer, whereas the shorter single-stranded TRS12 occupies two sites per dimer. This is consistent with the surface analysis described in next section, in which the CTD dimer has one central positively charged groove along the dimerisation interface and two symmetry-related peripheral basic patches, one on each protomer. For the NTD, all three RNAs gave n values close to one (0.8–0.9), as expected for a monomeric domain binding a single RNA molecule.

### The N protein CTDs of different coronaviruses contain a positive surface

We performed multiple sequence alignment of the CTD region of N proteins from different coronaviruses to understand the degree of sequence conservation among them. As shown in Supplementary Figure S2A, the SARS-CoV-2 N protein CTD expectedly shares the highest sequence similarity with SARS-CoV N protein CTD, exhibiting an identity of 95.76%. On the other hand, SARS-CoV-2 N protein CTD shared lower sequence identities with other coronaviruses, including MERS-CoV (55.08%), MHV (36.52%), and HCoV-NL63 (29.73%).

Despite the differences in sequence conservation, the overall three-dimensional structures of different coronaviruses N protein CTDs are highly similar (Supplementary Figure S2B). Structural comparison revealed the presence of a conserved positively charged groove on the α-helical face of the N protein CTD dimer in all the proteins, formed by the clustering of positively charged residues such as K256, K257, K261, and R262 in the SARS-CoV-2 N-CTD. This conserved electropositive surface was proposed to play an important role in RNA interaction in several previous studies ^22,23,38,48–50^.

The comparison of the electrostatic surface potentials of the opposite β-sheet face of the structures also showed some differences among different proteins. The MERS-CoV N protein CTD showed a positive charge in central region, whereas both SARS-CoV-2 and SARS-CoV N proteins CTDs have a negatively charged surface in the corresponding region. The β-sheet face of HCoV-NL63 contains more negatively charged central patch. These differences in surface electrostatics may contribute to variations in RNA recognition and binding modes among coronavirus N protein CTDs.

There is another positively charged surface on each of the protomer of the CTD dimer, comprising of positively charge residues Lys256, Arg259, and Arg262 of one monomer as well as Lys338 of another monomer. This site was previously identified to bind the ATP molecules ^39^.

### NMR titrations reveal dynamic CTD-TRS RNA Interactions

#### CTD - TRS23 titration

To probe the interaction between the CTD and TRS23 RNA, we performed NMR titrations experiments monitoring imino proton resonances of TRS23 RNA upon addition of increasing concentration of CTD. The imino resonances showed peak broadening, and at approximately two equivalents of CTD, the peaks were completely broadened indicating the binding between RNA and protein in the intermediate exchange timescale (Supplementary Figure S3).

We next followed the interaction between CTD and TRS23 by monitoring the changes in the ^1^H-^15^N HSQC spectra of CTD upon titration of TRS23 RNA. Throughout the NMR titrations, molar ratios are given as CTD monomer:RNA. Because CTD is dimeric, a ratio of 1:1 corresponds to two RNA molecules per CTD dimer. Addition of RNA resulted in severe peaks broadening and decrease in the peak intensities in the ^1^H-^15^N HSQC spectra of CTD (Figure 3A). Even at very low protein to RNA ratio of 1:0.25, we observed significant peak broadening and the loss of peak intensities for the majority of the residues. At 1:1 protein to RNA ratio, nearly all CTD peaks disappeared from the spectrum. This behavior suggests that the observed line broadening is likely arising from at least two contributing processes: (i) formation of large oligomeric CTD-TRS23 assemblies which result in slow rotational motions and consequently leading to the broadening of peaks in the ^1^H–^15^N HSQC spectrum, and/or (ii) binding-induced chemical exchange in the intermediate exchange timescale.

**Figure 3.**
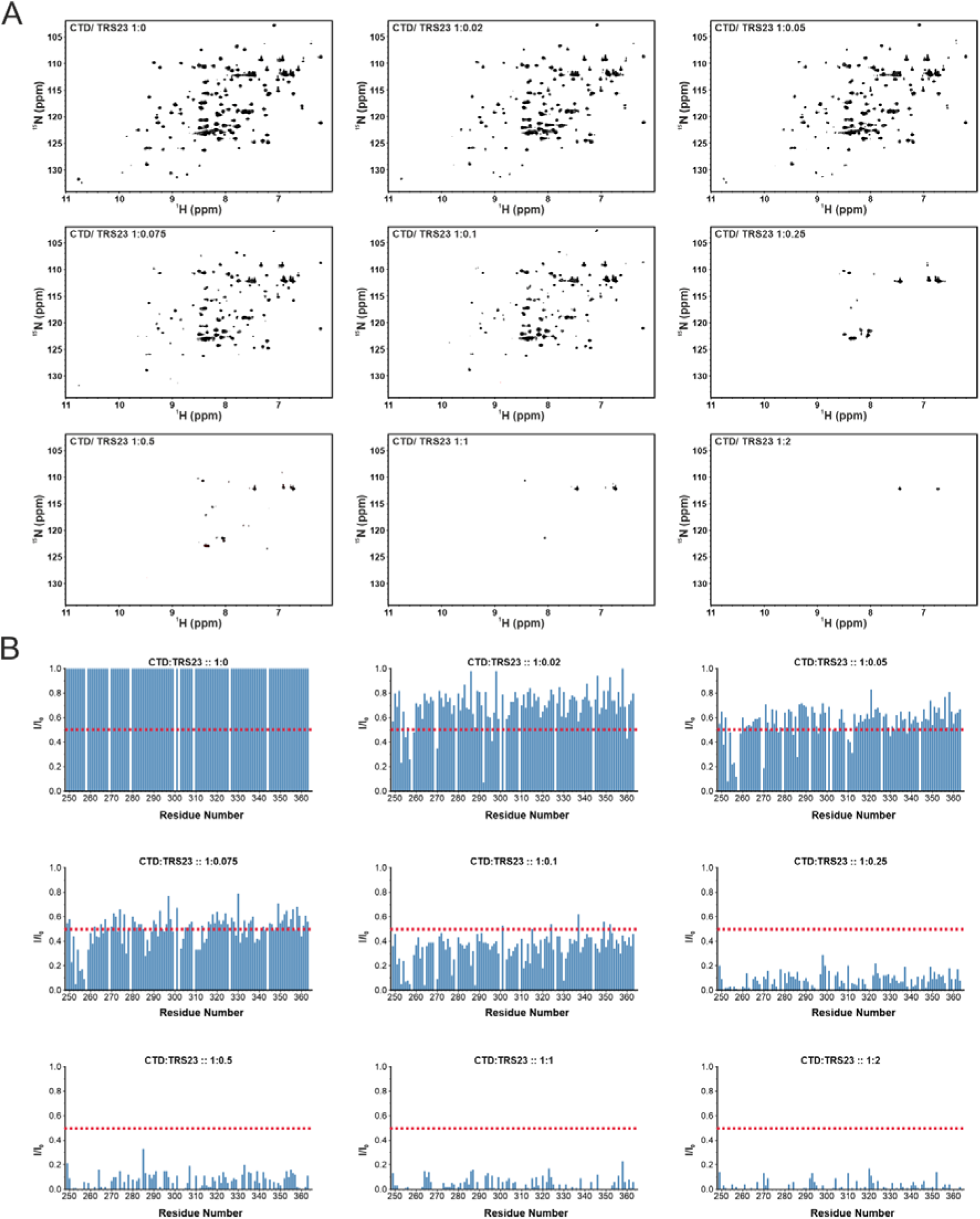
NMR titration of CTD with stem-loop TRS23 RNA. **(A)** Series of 2D ^1^H-^15^N HSQC spectra of ^15^N-labeled CTD in complex with TRS23 RNA. The molar ratio of CTD:TRS23 (1:0, 1:0.02, 1:0.05, 1:0.075, 1:0.1, 1:0.25, 1:0.5, 1:1, and 1:2) at each titration point is mentioned. Addition of increasing amount of TRS23 resulted in severe line broadening leading to gradual reduction and disappearance of CTD resonances. **(B)** Quantitative analysis of peak intensity changes during the course of CTD-TRS23 titration. For each titration point, peak intensities were normalized against the free CTD and plotted as the normalised intensity ratio (*I/I_₀_*) vs CTD residues, where *I* represents the peak intensity at a given protein:RNA ratio and *I_₀_* corresponds to the peak intensity in the absence of RNA. The red dashed line indicates an I/I*_₀_* value of 0.5, corresponding to a 50% reduction in signal intensity.

Severe line broadening of the resonances without any perturbations in the chemical shifts, hampered our efforts to perform CSP mapping to identify the RNA-binding interface on the protein. We undertook an alternate approach to characterise this interaction. We quantified the decrease in peak intensities of residues at each titration point by calculating the peak intensity ratios (*I/I_0_*), where I and I_0_ represent the peak intensities in the presence and absence of RNA, respectively. These intensity plots were plotted for each titration point (Figure 3B).

Residues that showed more than 50% loss in the signal intensity (I/I_0_ < 0.5) at each titration point were mapped onto the surface of the CTD structure (Figure 4). A previously determined structure of SARS-CoV-2 N protein CTD (PDB ID 6YUN) was used for this analysis ^51^. This analysis showed a gradual loss of peak intensities of the CTD residues upon addition of stem-loop TRS23 RNA. At the early points of titrations, intensities of residues in and around the positive grove of the CTD lost the intensities, however at the ratio of 1:0.1 CTD to TRS23 RNA, nearly all the residues showed drastic reduction in the peak intensities (Figure 4). These results suggest that the initial contact or binding point for TRS23 RNA might be the positively charged surface (formed by the dimerization interface on the a-helical side) of the CTD. However, with the increasing amount, TRS23 RNA is likely binding to other places on CTD leading to the formation of higher molecular weight complexes.

**Figure 4.**
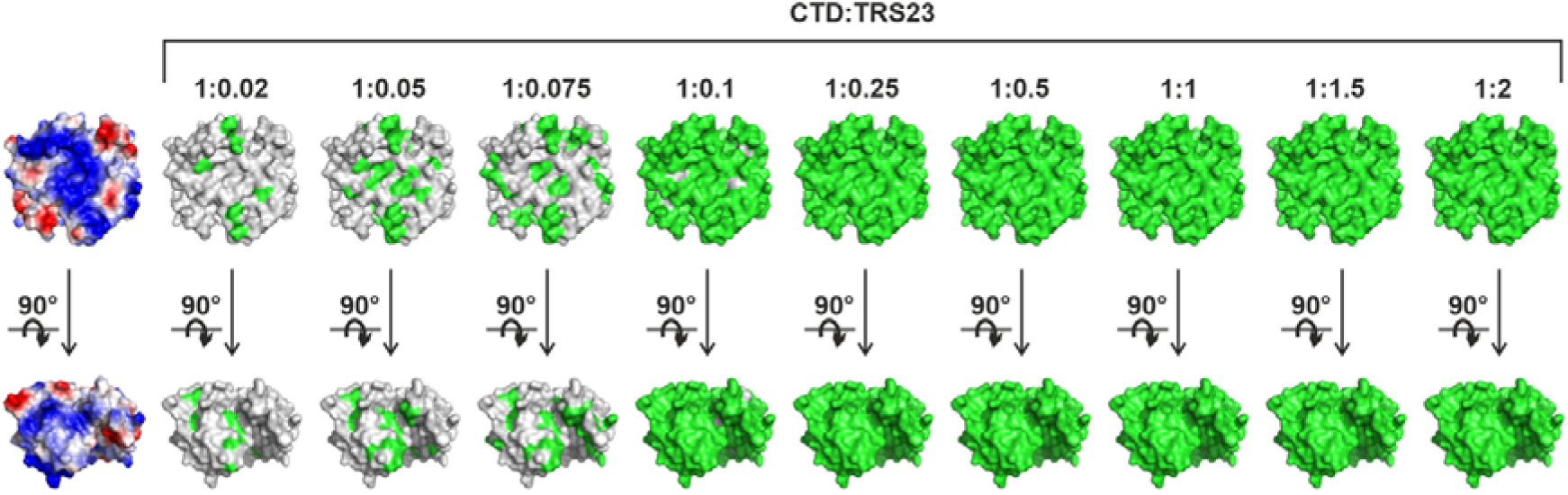
Structural mapping of residues showing 50% peak intensity reduction during CTD–TRS23 titration. Electrostatic surface potential of CTD structure (taken from PDB ID 6YUN) is shown in the left most panel for the reference, showing the distribution of positively (blue) and negatively (red) charged regions. Residues with normalized peak intensities (*I/I_₀_*) below 0.5 at each CTD:TRS23 molar ratio (1:0.02, 1:0.05, 1:0.075, 1:0.1, 1:0.25, 1:0.5, 1:1, 1:1.5, and 1:2) were mapped onto the CTD structure and are highlighted in green. The lower panels represent the same structures rotated by 90° about the x-axis to provide an alternative view of the spatial distribution of the affected residues.

#### CTD – TRS12 titration

In the next set of experiments, CTD was titrated with increasing concentration of single-stranded TRS12 RNA. The titration was followed by monitoring the 2D ^1^H–^15^N HSQC spectra of CTD at each titration point. In case of TRS12 titration also we observed peak broadening in the CTD ^1^H–^15^N HSQC spectra indicative of binding in the intermediate exchange regime (Figure 5A). However, *I/I_₀_* analyses for TRS12 titrations showed a less attenuated loss of signal intensity at each titration point compared to the TRS23 titration (Figure 5B).

**Figure 5.**
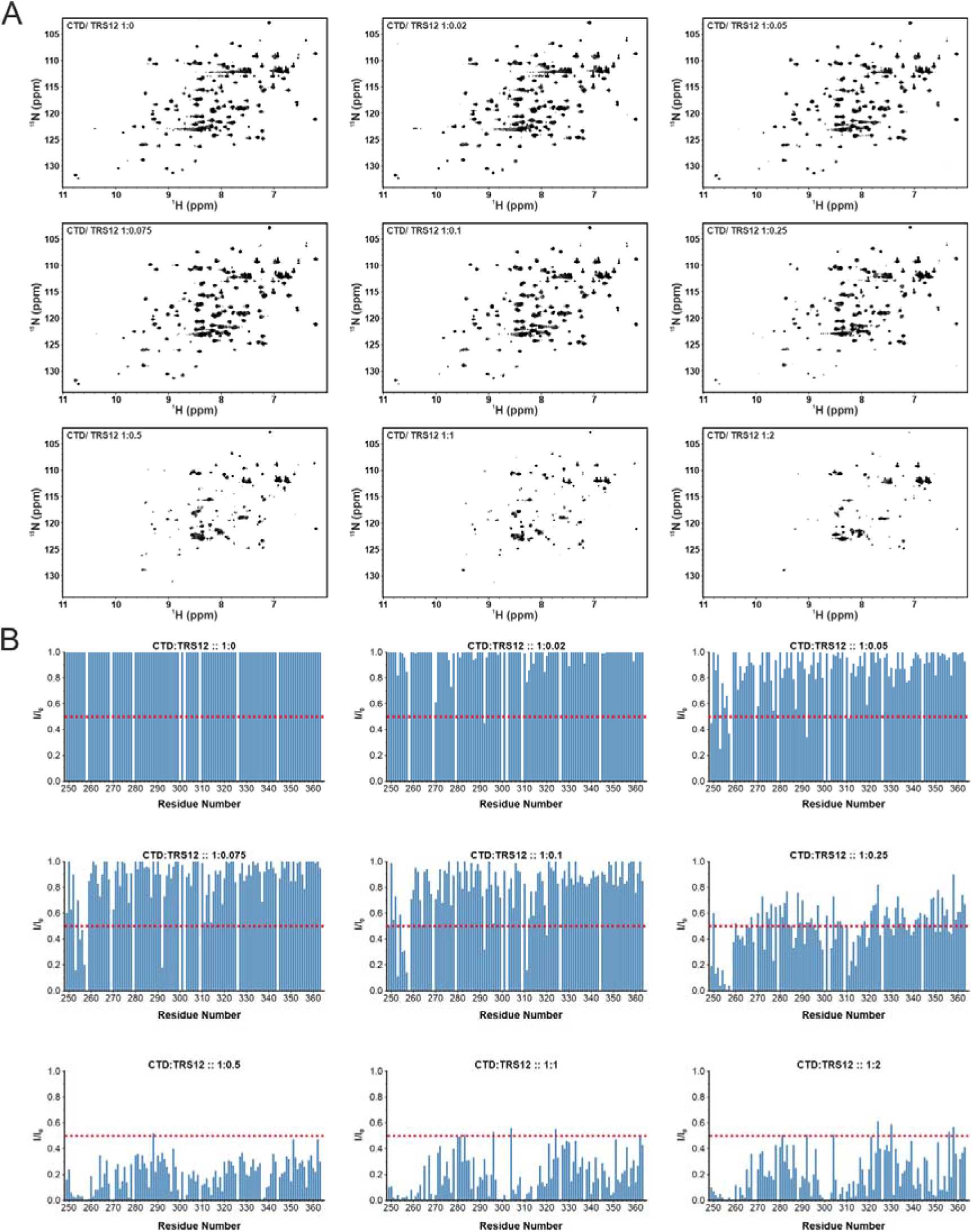
NMR titration of CTD with single-stranded TRS12 RNA. **(A)** Series of 2D ^1^H-^15^N HSQC spectra of ^15^N-labeled CTD in complex with TRS12 RNA. The molar ratio of CTD:TRS12 (1:0, 1:0.02, 1:0.05, 1:0.075, 1:0.1, 1:0.25, 1:0.5, 1:1, and 1:2) at each titration point is mentioned. Addition of increasing amount of TRS12 resulted in severe line broadening leading to gradual reduction and disappearance of several CTD resonances. **(B)** Quantitative analysis of peak intensity changes during the course of CTD-TRS12 titration. For each titration point, peak intensities were normalized against the free CTD and plotted as the normalised intensity ratio (*I/I_₀_*) vs CTD residues, where *I* represents the peak intensity at a given protein:RNA ratio and *I_₀_* corresponds to the peak intensity in the absence of RNA. The red dashed line indicates an I/I_₀_ value of 0.5, corresponding to a 50% reduction in signal intensity.

Residues that showed more than 50% loss in the signal intensity (*I/I_0_* < 0.5) at each titration point were mapped onto the surface of the CTD structure (Figure 6). A previously determined structure of SARS-CoV-2 N protein CTD (PDB ID 6YUN) was used for this analysis. At the early points of titrations, some of the residues at the periphery of the CTD lost their intensities, however gradually, several residues spread throughout the CTD showed decrease in the intensity (Figure 6). These results suggest that the initial contact or binding point for singles-stranded TRS12 RNA might be the positively charged surface located at the periphery on each protomer of CTD in CTD dimer. However, with the increasing amount of RNA, TRS12 (as in the case of TRS23) is likely binding to other places on CTD leading to formation of higher molecular weight complexes.

**Figure 6.**
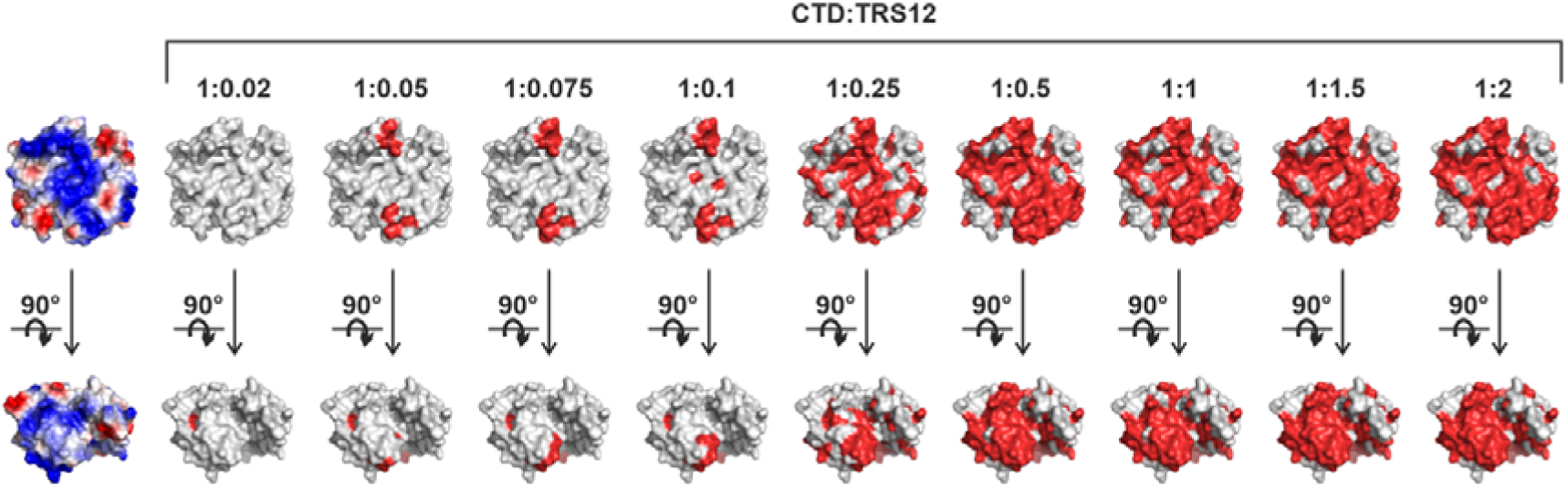
Structural mapping of residues showing 50% peak intensity reduction during CTD-TRS12 titration. Electrostatic surface potential of CTD structure (taken from PDB ID 6YUN) is shown in the left most panel for the reference, showing the distribution of positively (blue) and negatively (red) charged regions. Residues with normalized peak intensities (*I/I_₀_*) below 0.5 at each CTD:TRS12 molar ratio (1:0.02, 1:0.05, 1:0.075, 1:0.1, 1:0.25, 1:0.5, 1:1, 1:1.5, and 1:2) were mapped onto the CTD structure and are highlighted in green. The lower panels represent the same structures rotated by 90° about the x-axis to provide an alternative view of the spatial distribution of the affected residues.

#### CTD – TRS7 titration

In the next set of experiments, the ^15^N-labelled CTD was titrated with single-stranded TRS7 RNA and 2D ^1^H–^15^N HSQC spectra was recorded at each step of titration (Figure 7A). In this case, we observed gradual shifts of several distinct peaks (rather than the disappearance) upon addition of RNA. This suggest that CTD-TRS7 interaction is in the fast exchange regime, indicating weak binding and may not be forming higher oligomeric assemblies. This allowed us to track the perturbed resonances in the ^1^H–^15^N HSQC spectra.

**Figure 7.**
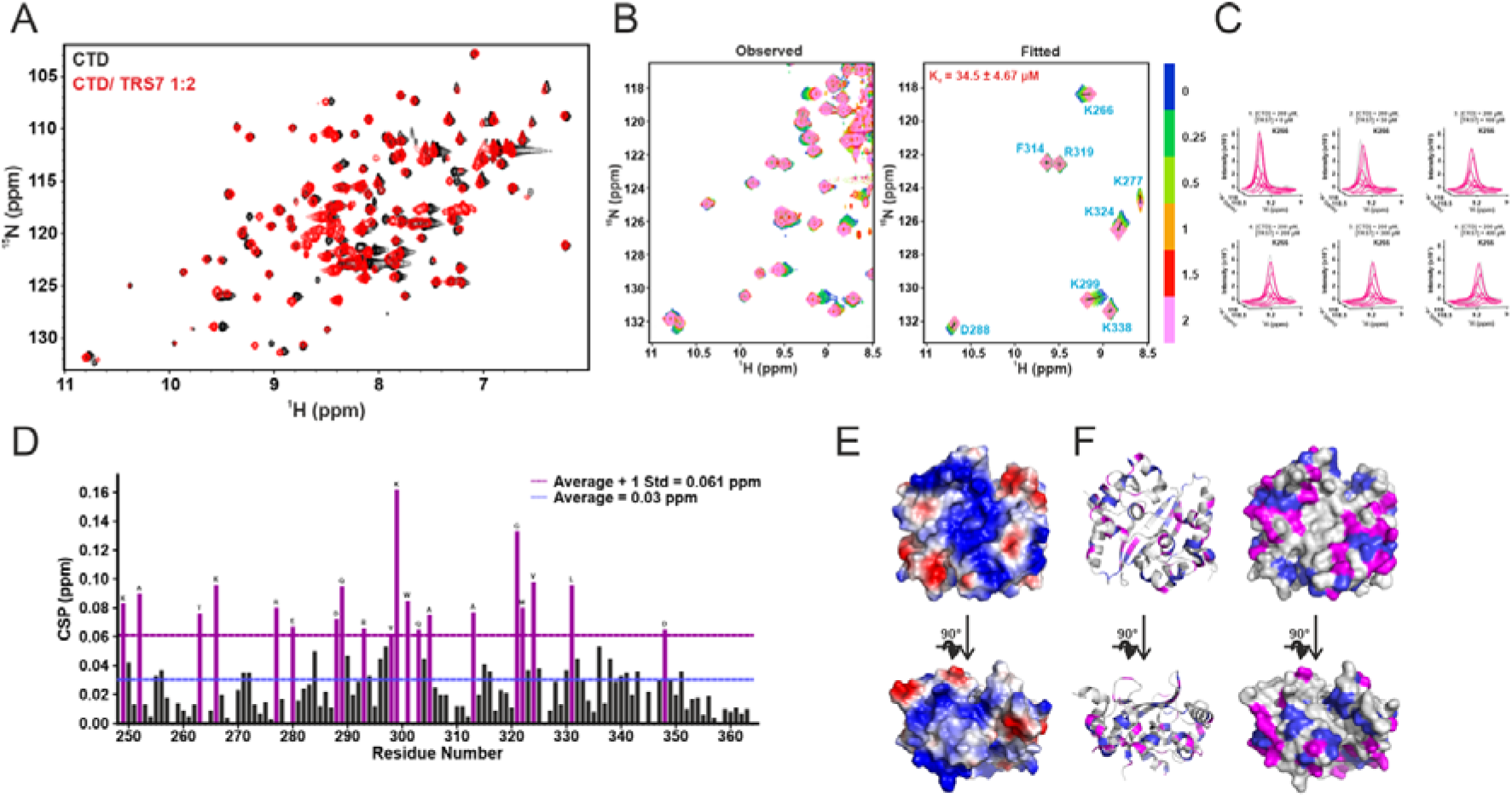
NMR titration of CTD with single-stranded TRS7 RNA. **(A)** Overlay of the 2D ^1^H-^15^N HSQC spectra of ^15^N-labeled CTD in the free state (black) and in the presence of TRS7 at a 1:2 molar ratio (red). **(B)** 2D line shape analysis of the interaction of CTD with TRS7 using NMR TITAN software. Representative regions of the experimental (2D ^1^H-^15^N HSQC) spectra used for TITAN analysis (left) and the corresponding spectra generated from the global fit (right). TITAN fitting of the NMR titration data yielded a K_d_ of 34.5 ± 4.67 _μ_M for the CTD-TRS7 interaction. **(C)** Three-dimensional overlay of observed (grey) and fitted (magenta) peak of a representative residue K266 to show quality of the fit. **(D)** Residue-wise CSP analysis of CTD upon TRS7 binding. CSP values were calculated from the overlaid ^1^H-^15^N HSQC spectra and plotted against the CTD residues. The blue line represents the average CSP value, while the magenta line indicates the average plus one standard deviation. Residues exhibiting CSP values above average deviation were considered significantly perturbed residues. **(E)** Electrostatic surface potential representation of CTD showing the distribution of positively charged (blue), negatively charged (red), and neutral (white) regions. The structure is displayed in three orientations to facilitate visualization of the surface properties. **(F)** Significantly perturbed residues with CSP values greater than the average plus one standard deviation are highlighted in magenta whereas residues with CSP values above average are shown in blue (ribbon form - left and surface form - right). The bottom panel corresponds to structure rotated by 90°, relative to the top panel.

To determine the binding affinity of CTD and TRS7 interaction we employed TITAN software which analyses the chemical shift perturbations as observed in the ^1^H–^15^N HSQC spectra of the in ^15^N-labelled CTD upon titration with TRS7 RNA (Figure 7B and 7C). The TITAN software performs two-dimensional lineshape analysis by fitting and simulating the ^1^H–^15^N HSQC spectra acquired across the titration series, thus enabling the determination of both the K_d_ and K_off_ values. We observed good agreement between the experimental and simulated lineshapes, which validated the quality of the fit and reliability of parameters derived from it. Our analysis yielded a K_d_ of 34.5 ± 4.67 μM and K_off_ of 5516.45 ± 1606.5 s^-1^. This indicates the binding of CTD with TRS7 is weaker under given experimental conditions.

The observed CSPs at saturating CTD to TRS7 RNA ratio of 1:2 were quantified and plotted against the residues of CTD (Materials and Methods) (Figure 7D). CTD residues (K249, A252, T263, K266, R277, E280, D288, Q289, R293, Y298, K299, W301, Q303, A305, A313, G321, M322, V324, L331, D348) that showed more than average and average plus one standard deviation CSPs upon TRS7 binding were classified as significant perturbations and mapped to the CTD structure. A previously determined structure (PDB ID 6YUN) and deposited chemical shifts (BMRB entry 50518) of CTD was used for the analysis. Interestingly, the residues that showed significant perturbation were distributed all over the structure including the residues around the positively charged basic patch of CTD (Figure 7E and 7F).

### CTD-TRS binding insights from AlphaFold3

To gain structural insights into the interaction of CTD with different TRS RNAs, structural models of CTD in complex with TRS7, TRS12, and TRS23 RNAs were generated using AlphaFold 3 (AF3) (Supplementary Figure S4A-C). The obtained structures are visualised and analysed in PyMol. In AF3 the predicted template modeling (pTM) and interface predicted template modeling (ipTM) scores greater than 0.8 corresponds to high-confidence predictions. The predicted CTD-RNA complexes generated in this study showed pTM and ipTM scores greater than 0.8, indicating high confidence in the predicted models.

The AF3 models predicted that all TRS RNAs associate with the positively charged surface of CTD, although distinct binding orientations were observed. While TRS7 and TRS12 were predicted to bind along the outer surface of the CTD dimer, TRS23 preferentially localized at the dimeric interface(Supplementary Figure S4A-C). Consistent with these observations, a 2:1 CTD:RNA stoichiometry was predicted for the CTD:TRS23 complexes, which is in agreement with the stoichiometry obtained from ITC measurements. In contrast, CTD:TRS7 and CTD:TRS12 complexes displayed a 1:1 binding stoichiometry.

Despite differences in the orientation and extent of RNA engagement, all predicted complexes localized the RNA molecules to positively charged patches on the CTD surface, supporting the electrostatic nature of CTD–RNA interactions.

### Size-exclusion chromatography reveals TRS RNA-induced assemblies of CTD

To further investigate the possible oligomerisation of CTD upon binding to TRS RNAs, we employed size-exclusion chromatography (SEC). CTD was incubated with TRS23, TRS12, and TRS7 RNA individually and the complexes were individually run on a size-exclusion chromatography (SEC) column (Material and Methods). The CTD alone and RNA alone samples were run separately as control. Free CTD eluted as a low-molecular-weight species consistent with its dimeric state (MWt ∼ 30.8 kDa) (Figure 8A). CTD-TRS23 complex eluted in multiple bands corresponding to higher molecular weight assemblies (Figure 8A). The major peak corresponded to a relative estimated molecular weight of around 208.8 kDa. Taking the mass of the CTD dimer as 30.8 kDa and of TRS23 as ∼7.4 kDa, this would correspond to a hetero-hexameric assembly of CTD-TRS23 (six CTD dimers with six TRS23 molecules). We note that apparent molecular weights derived from elution volume assume globular shape, and that an extended or asymmetric protein-RNA assembly will elute earlier than its true mass. CTD-TRS12 complex eluted at higher molecular weight assembly corresponding to an approximate molecular weight of 36kDa, suggesting an assembly of one CTD dimer and two TRS12 RNAs (Figure 8B). However, we also observed smaller peak corresponding to higher molecular weight assembly of CTD-TRS12 RNA (Figure 8B).

**Figure 8.**
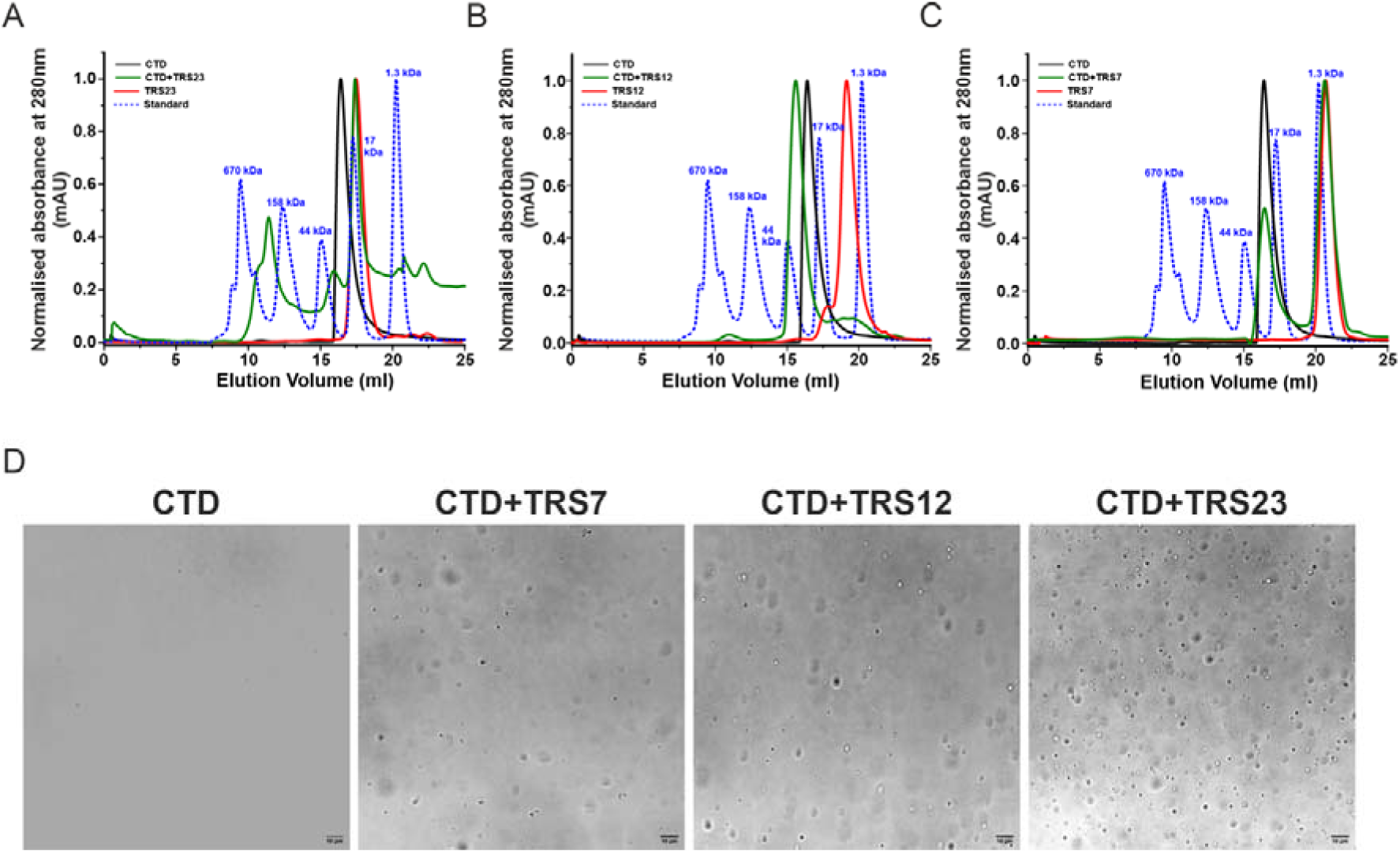
Analytical size-exclusion chromatography (SEC) and phase separation analyses of CTD in the presence of TRS RNAs. **(A–C)** Analytical SEC profiles of CTD, individual TRS RNAs, and their complexes on a Superdex 200 column. Elution profiles of free CTD (black), free RNA (red), and the corresponding CTD–RNA complexes (green) are shown along with molecular weight standards (blue dashed line). CTD in complex with stem-loop TRS23 RNA **(A)**, CTD in complex with single-stranded TRS12 RNA **(B)**, and CTD in complex with single-stranded TRS7 **(C)**. Shifts towards higher molecular weight in the SEC elution profiles relative to their free components shows the formation of CTD–RNA complexes. The elution positions of molecular weight standards are shown for the reference. **(D)** Representative differential interference contrast (DIC) images showing phase separation of CTD in the absence and presence of TRS RNAs at a 1:1 molar ratio. Under the experimental conditions CTD alone does not exhibit detectable condensates formation. However, the addition of TRS7, TRS12, or TRS23 promote the formation of liquid-like condensates, indicating that RNA binding enhances the phase separation propensity of CTD. Scale bars, 10 μm. Maximum number of droplets were observed in case of TRS23 RNA.

The CTD-TRS7 complex eluted at a column volume corresponding to an approximate molecular of 33.4 kDa (Figure 8C). This would correspond to an assembly of one CTD dimer and two TRS7 RNAs. Taken together, SEC results showed that CTD forms an higher molecular weight assembly with TRS23 and TRS12 RNAs, which is in agreement with the NMR titration results.

### N protein CTD undergoes phase-separation in the presence of TRS RNAs

N protein has been shown to undergo liquid-liquid phase separation (LLPS). Both folded and disordered regions of N protein contribute it its phase-separation, with the disordered region being the main contributor. Lkr region of N protein has been shown to undergo spontaneous phase separation in vitro. However, CTD alone has not been shown to undergo phase separation. As we have shown that CTD form higher oligomeric complexes upon TRS RNA binding, we next checked if it can also induce its phase separation of CTD *in vitro*.

The ability of the CTD to undergo RNA-induced phase separation was examined using differential interference contrast (DIC) microscopy. As expected CTD alone did not undergo phase separation under the tested experimental conditions tested (Figure 8D). Interestingly, upon addition of TRS7, TRS12, and TRS23 RNAs, the protein solution became visibly turbid (Figure 8D). Under the DIC microscopy, we observed micron-sized droplets characteristic of liquid–liquid phase separation. Increasing the RNA concentration led to a progressive increase in both the number and apparent density of droplets in all the cases. However, TRS23 induced most robust droplet formation compared with TRS7 and TRS12 RNAs under same set of conditions (Figure 8D). These results qualitatively show that CTD forms robust droplets in the presence of stem-loop TRS23 RNA.

## Discussion

Modular SARS-CoV-2 nucleocapsid protein has been shown to contain multiple RNA binding sites along its sequence. The N-terminal domain (NTD) was the first domain that was found to have RNA binding capacity ^52,53^. Later, the C-terminal dimerization domain (CTD) and the different disordered region of the N protein were also shown to bind RNA ^13,20,23,54,55^. However, the molecular details of the RNA interaction and sequence/structural preference of the CTD and other regions is not clearly understood.

The CTD, besides being responsible for the dimerization of the N protein, is also required for the higher-order assembly of the viral ribonucleoprotein complex ^23^. Past studies have shown that the CTD is not merely a passive oligomerization scaffold but also possesses intrinsic RNA binding activity, which is independent of the RNA binding NTD. Isolated CTD forms a stable dimer in solution and is capable of directly binding RNA, thereby suggesting that RNA recognition and self-association maybe functionally coupled within this domain. Crystal structures of the SARS-CoV-2 CTD had revealed a tightly associated dimer with a highly electropositive surface along the dimeric interface on the alpha-helical face of the CTD dimer ^51^. Subsequently, several studies had proposed that this positively charged grooves on the dimer interface could serve as RNA-binding sites ^22,23,38,48–50^.

In this study, we have attempted to delineate the RNA binding mechanism of N protein CTD by systematically studying its interaction with the RNA sequences from the transcriptional regulatory sequence (TRS) of SARS-CoV-2 genome. The SARS-CoV-2 N protein’s CTD was previously shown to interact with the central TRS sequence (5’-ACGAAC-3’) ^27^. We designed three RNAs that contain the TRS sequence element: stem-loop TRS23, single-stranded TRS12, and single-stranded TRS7 RNAs. The results from isothermal titration calorimetry titrations showed that CTD interacts with all the three RNAs, however interestingly, we found that stem-loop TRS23 binds CTD with better affinity than the single stranded TRS12 (∼6.4-fold higher affinity) and TRS7 (∼19.6-fold higher affinity) RNAs. This showed that CTD has preference for the stem-loop RNA than the single stranded RNA. On the other hand, ITC results showed that the NTD binds single-stranded RNA with higher affinity than the stem-loop RNA. The results also revealed that the CTD binds stem-loop TRS23 RNA with more than six-times better affinity than the NTD, clearly suggesting that CTD has preference for the stem-loop RNA.

The NMR titration data provide additional insight into the nature of the CTD-RNA interaction. In our experiments, addition of TRS23 to the CTD caused severe peak broadening and near-complete disappearance of resonances in the ^1^H–^15^N HSQC spectra even at relatively low RNA:protein ratios suggesting formation of a larger assembly of protein-RNA, with an interaction in the intermediate-exchange time regime on NMR time scale. Similar observation was made for CTD-TRS12 titration; however, the line broadening was less severe than the CTD-TRS23 titration at each titration point. In contrast, titration with TRS7 produced gradual chemical shift changes in the CTD ^1^H–^15^N HSQC spectra characteristic of a weaker and faster-exchanging interaction. These contrasting spectral responses suggest that structured TRS RNA does not merely bind the CTD more tightly but also promotes formation of larger or more slowly tumbling assemblies. This observation was further supported by our analytical size-exclusion chromatography (SEC) experiments. SEC elution profile of CTD-RNA complexes clearly showed formation of higher molecular weight aggregates in CTD-TRS23 complex. CTD-TR12 complex also showed low level of higher molecular weight aggregates compared to the CTD-TRS23 complex. On the other hand, the CTD-TRS7 complex eluted at an apparent mass of 33.4 kDa, only marginally larger than the free CTD dimer, giving no evidence for higher-order assembly, matching with the results of the NMR titrations. Taken together, these data argue that the CTD can switch from a soluble dimeric RNA-binding module to a higher-order assembly state depending on the RNA ligand.

Analysis of CTD-TRS23 and CTD-TRS12 NMR titrations revealed selective and differential decrease in the peak intensities of the CTD resonances in the HSQC spectra. This analysis revealed that in case of CTD-TRS23 titration, several peaks from the α-helical side of the dimerization interface start to disappear right from the very early step of titration (starting with 1:0.02 CTD:TRS23 RNA ratio). This may suggest that initially, TRS23 contacted this interface of the CTD. At the 1:0.25 and above CTD:TRS23 RNA ratio, almost all the CTD NH resonances in the ^1^H–^15^N HSQC spectra disappear or lost intensity more than 50% of the free CTD intensity. In case of CTD-TRS12 RNA titration, initially, we observed disappearance or loss of intensity (> 50%) of the peaks at the periphery or edge of the dimerization interface. Interestingly, in a previous study, these residues were found to be perturbed upon binding with the ATP ^39^. This suggest that these residues may constitute the first contact point for the TRS12 RNA on CTD. At the higher ratio of CTD to RNA, we observed further decrease in the peak intensity, however compared to TRS23, several peaks remained visible in the spectra (*albeit* with the reduced intensity).

NMR titrations results revealed that CTD binds TRS7 RNA with weaker affinity, which resulted in binding in fast-exchange timescale, allowing tracking of chemical shift perturbation. This analysis revealed that the residues which were perturbed significantly (CSP greater than average perturbation) were distributed all over the CTD structure. Taken together, all these observations, suggest that the structured stem-loop TRS23 RNA initially occupy the central positively charged groove on the dimeric interface of the CTD. Further addition of TRS23 drive formation of the higher oligomeric CTD-TRS23 complex. On the other hand, in case of single stranded RNAs (TRS12 and TRS7) multiple surface residues appear to contribute cooperatively rather than through a single well-defined binding pocket.

SARS-CoV-2 nucleocapsid protein has been well documented to undergo phase separation in vitro ^18,20,21,56,57^. Intrinsically disordered (IDR) regions flanking the ordered NTD and CTD are shown to be important for the spontaneous phase separation of the protein, even without RNA. Our observations on CTD interaction with RNA in this study led us to investigate if CTD alone or in the presence of RNA can undergoes in vitro phase separation. Our results showed that while CTD alone does not phase separate, in the presence of all the three TRS RNAs used in this study, it underwent phase separation. However, here again, the most robust phase separation was observed with the stem-loop TRS23 RNA. These observations are consistent with the previous studies, that had shown that not just RNA sequence or length but also structure acts as a key determinant of phase separation of the N protein of SARS-CoV-2. Structured or double-stranded RNA can enhance N protein condensate formation by increasing binding valency which aligns well with our CTD-TRS23 interaction observed in this study ^20,31^

In summary, the present study supports an expanded functional role for the SARS-CoV-2 N-protein CTD in viral RNA recognition. Together with the prior studies, our biophysical and NMR our data indicate that the CTD is a *bona fide* RNA-binding domain whose interactions depend strongly on RNA structure and are closely linked to higher-order assembly. The preferential recognition of stem–loop TRS RNA, the induction of oligomeric complexes, and the ability to drive phase separation together suggest that the CTD may contribute directly to spatial and temporal organization of viral ribonucleoprotein complexes during transcription and genome packaging. These findings strengthen the view that the CTD is not only central to N-protein self-association but also an important determinant of RNA-selective assembly in the coronavirus life cycle.

## Associated content

### Supplementary Material Description

Supporting information is provided in the SI file. Supporting Information includes Table T1 and Figures S1–S4

### Author contributions

M.S. conceived the research and acquired the funds. P.G. designed and carried out the experimental design and performed all the experiments. The manuscript was written by M.S. and P.G.

## Acknowledgments

The authors acknowledge funding for infrastructural support from the following programs of the Government of India: DST FIST, DST, DBT SAIF, UGC CAS, and the DBT-IISc partnership program. We acknowledge the divisional bioimaging facility for imaging. P.G. was supported by a Ph.D. fellowship from Ministry of Education (MoE).

Authors acknowledge funding support from the Anusandhan National Research Foundation (ANRF), Department of Science and Technology (DST), India (STAR award number STR/2021/000015 and CRG grant number CRG/2021/003597 to M.S.) and Ministry of Education (MoE). Research in M.S. lab is currently supported funding support from the Department of Biotechnology (DBT), India (grant number BT/PR56485/BMS/85/598/2024); the Anusandhan National Research Foundation (ANRF) (grant number ANRF/ARG/2025/007912/LS), and Indian Council of Medical Research (ICMR) (grant number IIRPSG-2025-01-01384).

## Abbreviations

SARS-Cov-2: severe acute respiratory syndrome coronavirus 2
LLPS: Liquid-liquid phase separation
NTD: N-terminal domain
CTD: C-terminal domain
N protein: Nucleocapsid protein
RNP: Ribonucleoprotein complex
LCD: Low complexity domain
IDR: intrinsically disordered region
ITC: isothermal titration calorimetry
TRS: transcriptional regulatory sequence

## Data availability

This study did not generate unique reagents. All the materials of this study are available from the lead contact upon reasonable request.

