## Supporting Information for "C-terminal domain of SARS-CoV-2 nucleocapsid protein binds transcriptional regulatory sequence RNA with high affinity and forms higher-order assemblies"

This file contains:

Supplementary Figures S1, S2, S3, and S4

Supplementary Table T1

**Supplementary Figure S1**

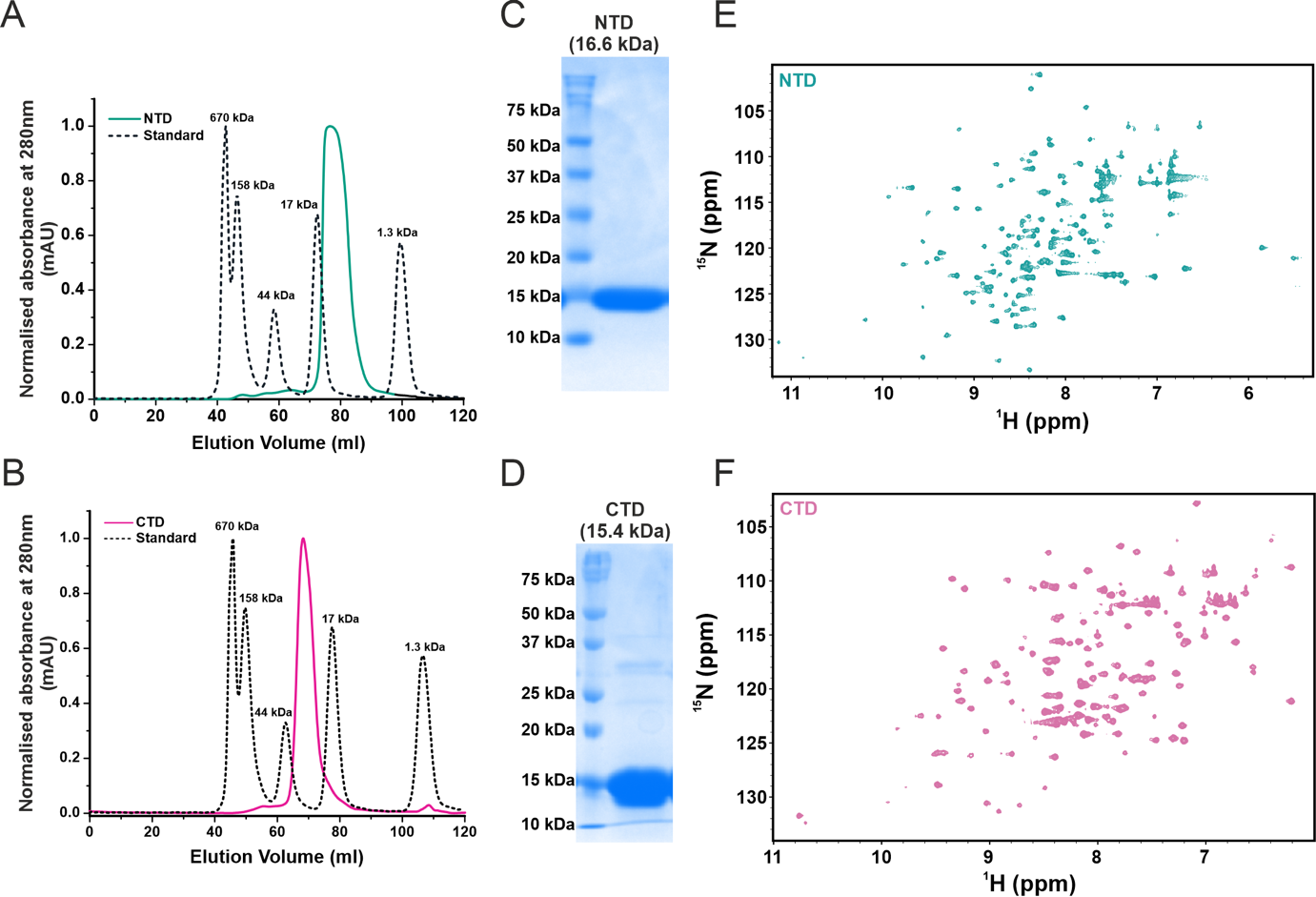

**Supplementary Figure S1.** Protein purification and biophysical characterization of N protein NTD and CTD. **(A)** Size-exclusion chromatography (SEC) profile of the NTD. The elution profile of NTD (cyan) is overlaid with molecular weight standards (black dashed line) for reference. The NTD elutes as monomer in solution. **(B)** SEC profile of the CTD. CTD elutes as a dimer in solution. The elution profile of CTD (magenta) is overlaid with molecular weight standards (black dashed line) used as reference. **(C)** SDS-PAGE analysis of purified NTD showing a single band corresponding to the expected molecular mass of ~16.6 kDa. **(E)** 2D ^1^H-^15^N HSQC spectrum of uniformly ^15^N -labelled NTD. The well-dispersed resonances indicate that NTD adopts a folded conformation in solution. **(D)** SDS–PAGE analysis of the purified CTD showing a band corresponding to the expected molecular mass of ~15.4 kDa. **(F)** 2D ^1^H-^15^N HSQC spectrum of uniformly ^15^N-labelled CTD recorded under the conditions described in the Materials and Methods. The observed chemical shifts are well dispersed, indicative of a well-folded protein in solution.

**Supplementary Figure S2**

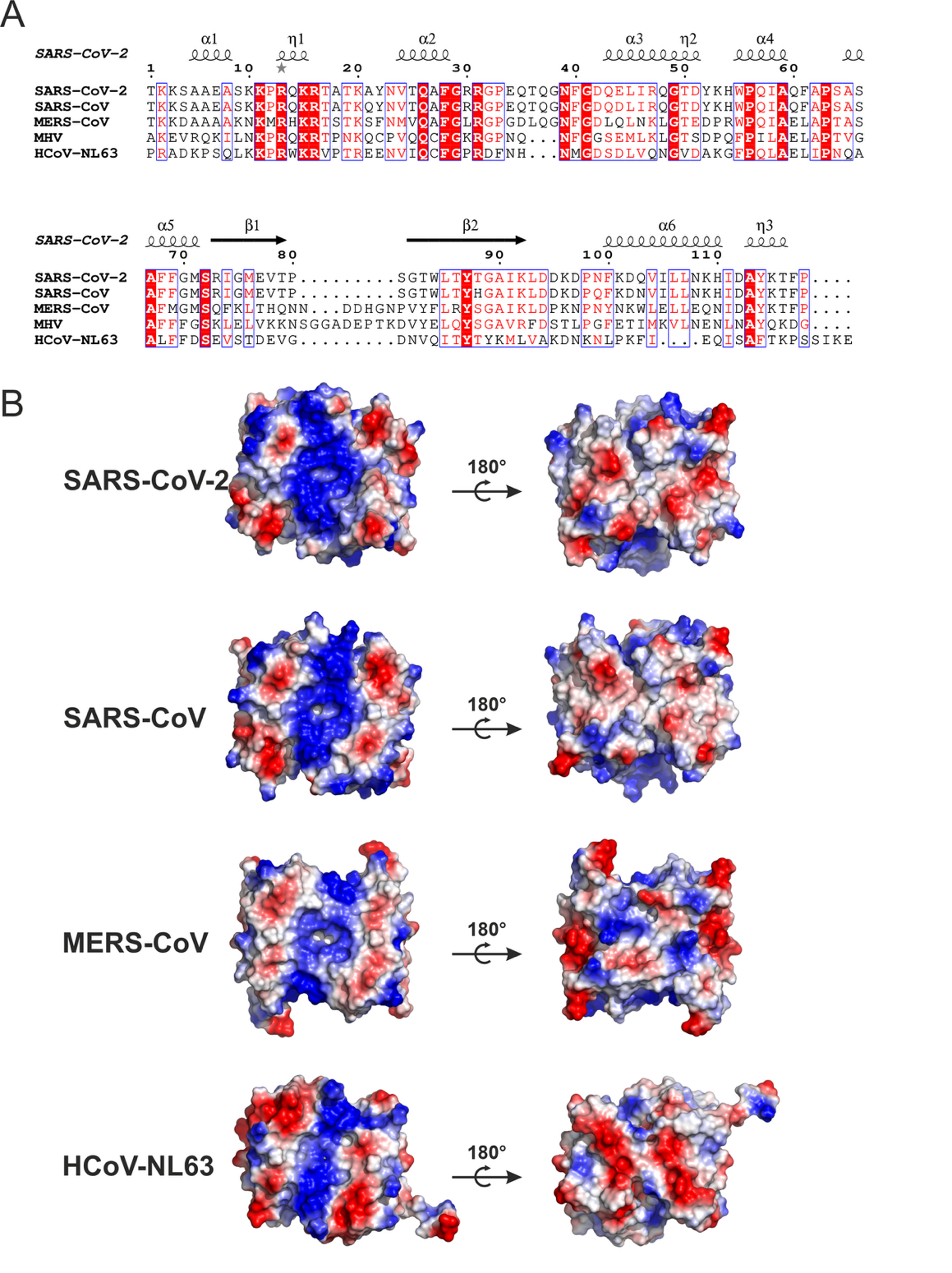

**Supplementary Figure S2.** Comparison of sequence and structures of CTD of coronaviruses. **(A)** Multiple sequence alignment of CTD from different coronaviruses. The secondary structure corresponding to the CTD is depicted at the top. The consensus sequence, including the conserved residues (red), is shown at the bottom. **(B)** Surface electrostatic potential of the CTDs from different coronaviruses. One side (consisting of alpha-helices) of the protein has positively charged surface (blue), while the other side (consisting of beta-sheet) has negatively charged surface (red).

**Supplementary Figure S3**

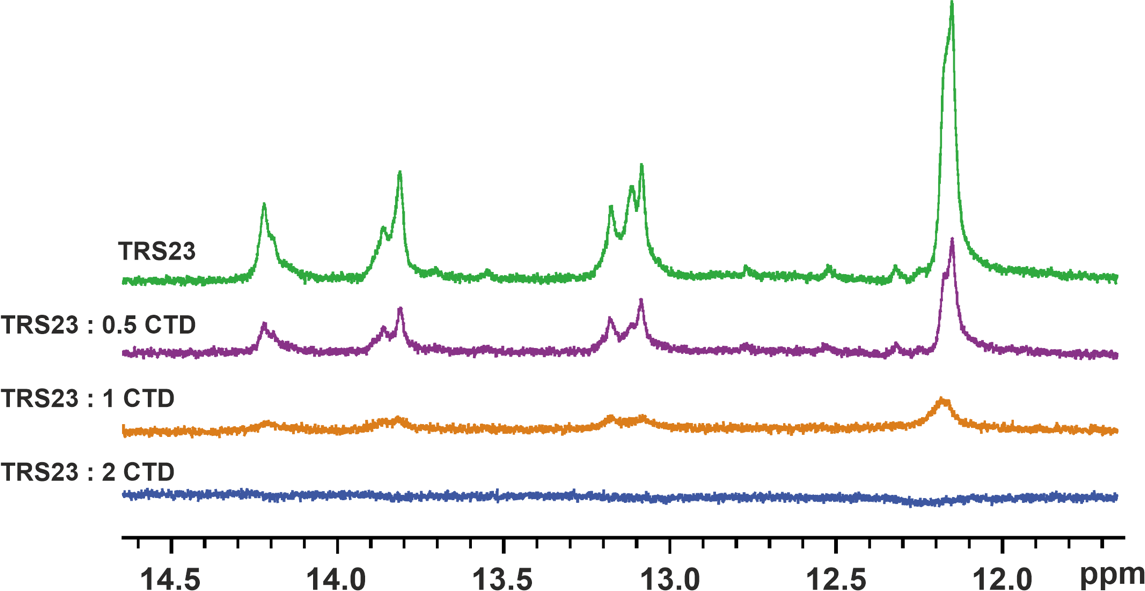

**Supplementary Figure S3. NMR titration of stem-loop TRS23 with CTD.** 1D ^1^H NMR spectra showing imino proton region of TRS23 RNA in the absence and presence of increasing concentrations of CTD. TRS23 forms a stem-loop structure containing four base pairs, giving rise to four distinct imino proton resonances in the spectrum of free RNA (green) and for TRS23:CTD molar ratios of 1:0.5 (purple), 1:1 (orange), and 1:2 (blue). Addition of CTD results in concentration dependent broadening and reduction in the intensity of TRS23 imino resonances, indicating an interaction between CTD and TRS23 RNA.

**Supplementary Figure S4**

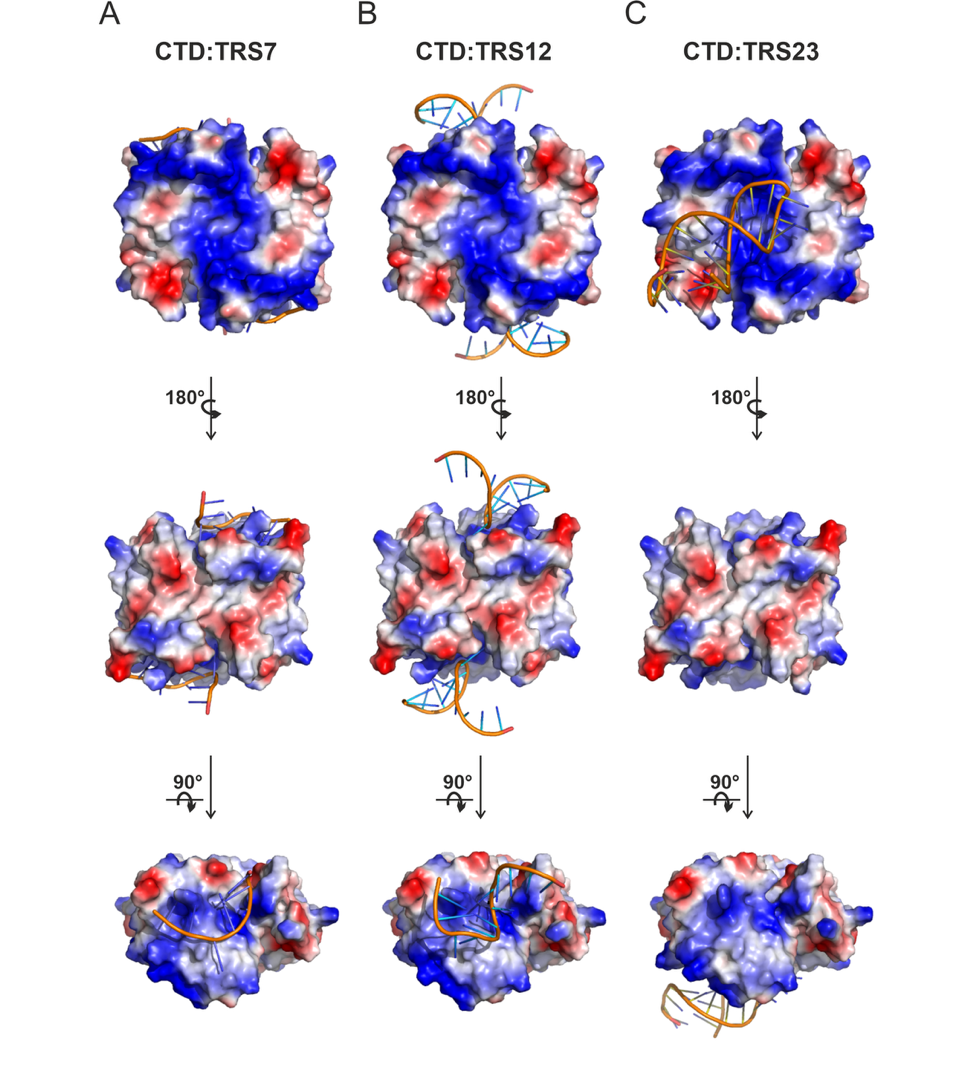

**Supplementary Figure S4.** AlphaFold3 predicted model structures of CTD in complex with different TRS RNA constructs. **(A)** CTD bound to single-stranded TRS7. **(B)** CTD bound to single-stranded TRS12, **(C)** CTD bound to stem-loop TRS23. CTD is shown as an electrostatic surface representation, with positively charged regions coloured blue, negatively charged regions coloured red, and neutral regions coloured white. RNA molecules are shown as cartoon representations. For each complex, the middle and bottom panel views are obtained after successive rotations of 180° and 90°, respectively, relative to the orientation shown in the top panel.

**Supplementary Table T1.** RNA sequences used in this study.

| **S. No.** | **RNA** | **Sequence** |
| --- | --- | --- |
| 1 | TRS23 | GGUGUUCUCUAAACGAACUUUAA |
| 2 | TRS12 | GGAAACGAACUU |
| 3 | TRS7 | ACGAACU |
